# Stepwise recombination suppression around the mating-type locus in the fungus *Schizothecium tetrasporum* (Ascomycota, Sordariales)

**DOI:** 10.1101/2022.07.20.500756

**Authors:** Nina Vittorelli, Alodie Snirc, Emilie Levert, Valérie Gautier, Christophe Lalanne, Elsa De Filippo, Ricardo C. Rodríguez de la Vega, Pierre Gladieux, Sonia Guillou, Yu Zhang, Sravanthi Tejomurthula, Igor V. Grigoriev, Robert Debuchy, Philippe Silar, Tatiana Giraud, Fanny E. Hartmann

## Abstract

Recombination is often suppressed at sex-determining loci in plants and animals, and at self-incompatibility or mating-type loci in plants and fungi. In fungal ascomycetes, recombination suppression around the mating-type locus is associated with pseudo-homothallism, *i*.*e*., the production of self-fertile dikaryotic sexual spores carrying the two opposite mating types. This has been well studied in two species complexes from different families of Sordariales: *Podospora anserina* and *Neurospora tetrasperma*. However, it is unclear whether this intriguing convergent association holds in other species. We show here that *Schizothecium tetrasporum*, a fungus from a third family in the order Sordariales, also produces mostly self-fertile dikaryotic spores carrying the two opposite mating types. This was due to a high frequency of second meiotic division segregation at the mating-type locus, indicating the occurrence of a single and systematic crossing-over event between the mating-type locus and the centromere, as in *P. anserina*. The mating-type locus has the typical Sordariales organization, plus a *MAT1-1-1* pseudogene in the *MAT1-2* haplotype. High-quality genome assemblies of opposite mating types and segregation analyses revealed a suppression of recombination in a region of 1.3 Mb around the mating-type locus. We detected three evolutionary strata, displaying a stepwise extension of recombination suppression, but no rearrangement or transposable element accumulation in the non-recombining region. Our findings indicate a convergent evolution of self-fertile dikaryotic sexual spores across multiple ascomycete fungi. The particular pattern of meiotic segregation at the mating-type locus was associated with recombination suppression around this locus, that had extended stepwise. This association is consistent with a recently proposed mechanism of deleterious allele sheltering through recombination suppression around a permanently heterozygous locus.

**AUTHOR SUMMARY:** Recombination allows faster adaptation and the purging of deleterious mutation but is often paradoxically lacking in sex chromosomes. It has been recently recognized that recombination can also be suppressed on fungal mating-type chromosomes, but the evolutionary explanation and the proximal mechanism of this phenomenon remain unclear. By studying here the sexual biology of a poorly studied mold living in rabbit dung, we reveal a striking convergence in three distant fungal lineages of an independently evolved association between the production of self-fertile sexual spores (carrying two nuclei with opposite mating types), a particular segregation of the mating-type locus and the lack of recombination on mating-type chromosomes, having evolved stepwise. Such a convergent association suggests causal relationships and will contribute to unveil the evolutionary causes of recombination suppression.

**Graphical summary:** 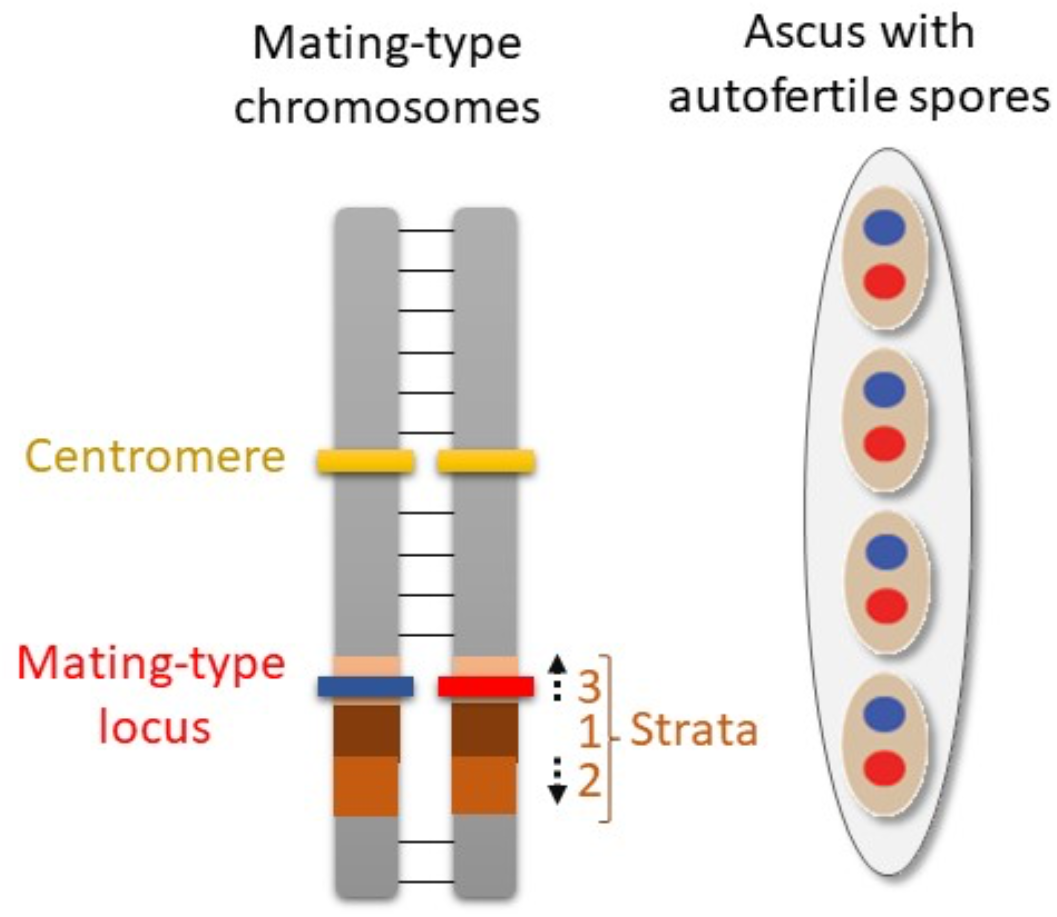

## INTRODUCTION

In eukaryotes, recombination is often considered to be a major long-term evolutionary advantage. Indeed, it prevents the accumulation of deleterious mutations (1) and facilitates the combination of multiple advantageous alleles in the same genome (2). However, convergent recombination suppression events have been observed at sex-determining loci in plants and animals, and at self-incompatibility or mating-type loci in plants and fungi (3). It has been suggested that this recombination suppression links lock-and-key genes together, enabling self-incompatibility systems to function correctly (4–6). Surprisingly, recombination suppression has often been found to extend distally, in a stepwise manner, from sex and mating-type loci, generating evolutionary strata of differentiation between haplotypes (7–10). The evolutionary causes of this stepwise recombination suppression beyond mating-compatibility loci remain a matter of debate (9–12). Sexual antagonism was long thought to be the main driving force generating evolutionary strata on sex chromosomes, through selection for the linkage to the sex-determining locus of genes with alleles beneficial in only one sex (12). However, it has been difficult to obtain definitive evidence supporting this hypothesis, and evolutionary strata have also been reported in fungi without sexual antagonism or other types of antagonism between mating types (8–10,13). Alternative hypotheses have been developed (10,11), based on associative overdominance associated with the sheltering of deleterious mutations (14), dosage compensation (15) or the spread of transposable elements (16).

Fungi are useful models for studies of the evolution of recombination suppression: they have many assets for evolutionary genomics studies (17) and several convergent recombination suppression events around mating-type loci have been observed in this group (8,10,18). Fungi have highly diverse modes of reproduction, mating systems (selfing versus outcrossing) and breeding systems (mating compatibility), with considerable differences sometimes reported between closely related species (19). Among all breeding systems, heterothallism was the most investigated and seems to display the most conserved scheme in fungi (20). In heterothallic fungi, mating can occur only between haploid cells carrying different alleles at the mating-type locus. The term “heterothallism” was initially coined after observations of *in vitro* crosses, in which mating could occur only between different thalli (*i*.*e*., mycelia or hyphal networks), each arising from a single spore. We now know that this heterothallism was due to the thalli mostly being haploid and carrying different alleles at the mating-type locus. Other fungi were described as “homothallic” because mating within a single thallus was possible. We now know that homothallism is possible because mating in these fungi does not need to occur between cells of different mating types. However, some fungi capable of displaying mating within a single thallus *in vitro* were later found to carry two types of nuclei, of different mating types, in each cell. They were therefore called pseudo-homothallic.

The pseudo-homothallic breeding system has been studied in detail in two species from different families of the order Sordariales (Ascomycota): *Neurospora tetrasperma* (Sordariaceae), which is typically found as orange blooms on scorched vegetation at burnt sites after a fire, and *Podospora anserina* (Podosporaceae), which is mostly found in herbivore dung. Sexual spore formation is similar in these two species (21). Karyogamy occurs between two haploid nuclei of different mating types, followed by meiosis, and then the post-meiotic mitosis of each meiosis product. The ascus produced therefore contains eight haploid nuclei, which are packaged in pairs to form four sexual spores (ascospores). The resulting ascospores are, thus, dikaryotic, *i*.*e*., each contains two haploid nuclei. A small fraction of asci (0.1% for *N. tetrasperma* and 1.0% in *P. anserina*) contains five ascospores, with one dikaryotic spore replaced by a pair of monokaryotic ascospores (*i*.*e*., ascospores carrying a single haploid nucleus), recognizable in *P. anserina* by their smaller size. In both species, the dikaryotic spores almost always contain nuclei of opposite mating types, and are therefore referred to as heterokaryotic ascospores, each of which germinates into a self-fertile mycelium (21). The particular pattern of segregation of the mating-type locus leads to the generation of nuclei of different mating types in close physical proximity, which can be compacted into a single ascospore. In *N. tetrasperma*, the mating-type locus segregates during the first meiotic division, due to the linkage between the centromere and the mating-type locus. During meiosis II, the spindles are parallel, and, following the post-meiotic mitosis, each ascospore is delineated around two non-sister nuclei, resulting in the compaction of opposite mating types in the same cell. In *P. anserina*, the mating-type locus segregates during the second meiotic division, due to the occurrence of a single and systematic crossing-over (CO) event between the centromere and the mating-type locus. During meiosis II, the spindles are aligned, leading to the same final nuclear distribution as in *N. tetrasperma*. After the post-meiotic mitosis, the ascospores formed contain nuclei of opposite mating types.

In these two pseudo-homothallic species, there are large non-recombining regions around the mating-type locus. In *N. tetrasperma*, the 7 Mb region without recombination encompasses the mating-type locus and the centromere (22). In *P. anserina*, a 0.8 Mb region devoid of recombination has been identified around the mating-type locus in the reference strain (23). Each of these two species belongs to a complex of cryptic species (22,24,25), which also all harbor a region of recombination suppression around their mating-type locus; however the size of the non-recombining region differs between species within each species complex (22,26,27). In both *N. tetrasperma* and *P. pseudocomata* (a cryptic species from the *P. anserina* species complex; (24)), the suppression of recombination has extended stepwise, generating evolutionary strata (10,22,26,27). It is intriguing that the presence of non-recombining regions around fungal mating-type loci and their stepwise extension seem to be associated with pseudo-homothallism in ascomycetes (10). One hypothesis that could account for this association is that most strains are heterokaryotic at the mating-type locus for most of their life cycle in pseudo-homothallic ascomycetes. This could allow recombination suppression to be selected as a means of sheltering recessive deleterious alleles, by linking them to a permanently heterozygous locus (14). Indeed, non-recombinant haplotypes with fewer recessive deleterious mutations than the population average should be selected for when rare, but if they increase in frequency, they will start to occur as homozygotes, exposing their recessive load, unless they capture a permanently heterozygous allele; in such cases, they continue to be selected and can rapidly reach their maximal frequency, *i*.*e*. becoming fully associated with the heterozygous allele close to which they appeared (14). The dikaryotic phase should also foster a stepwise extension of recombination suppression around the mating-type locus by the same mechanism (14). However, studies of additional pseudo-homothallic fungi are required, to assess the strength of the association between pseudo-homothallism and stepwise recombination suppression around the mating-type locus.

We therefore studied *Schizothecium tetrasporum* (previously known as *Podospora tetraspora* or *P. minuta var. tetraspora)*, a third pseudo-homothallic species from order Sordariales (21,28). *Schizothecium tetrasporum* belongs to the family Schizotheciaceae (also known as Neoschizotheciaceae), while *P. anserina* belongs to the Podosporaceae and *N. tetrasperma* to the Sordariaceae (29,30). The phylogenetic relationships between these three families are not well established, but they probably diverged over 150 million years ago (31). For example, the phylogenetic distance between *P. anserina* and *N. crassa*, a member of the Sordariaceae (like *N. tetrasperma)*, corresponds roughly to that between humans and fishes (32). Phylogenetically, *Schizothecium tetrasporum* is roughly equidistant from these two species (29). However, it displays striking cytological similarities to *P. anserina*, particularly in terms of dikaryotic spore production: in the FGSC7436 strain, 98% of asci were reported to produce four dikaryotic spores, with the spindles aligned in the ascus during meiosis II (21). However, this previous study did not clearly investigate whether the two nuclei of a dikaryotic spore contained opposite mating types and did not study the mating-type locus or recombination suppression.

We identified media suitable for reproducible and fast growth, and media for sexual reproduction in *S. tetrasporum*. We studied the dikaryotic strain CBS815.71, estimating the frequency of second-division segregation of the mating-type locus at 99.5%. This indicates the existence of a single and systematic CO event between the mating-type locus and the centromere. We also built high-quality genome assemblies of homokaryons (*i*.e., with only one nucleus type) with opposite mating types, making it possible: i) to elucidate the genetic organization of the mating-type locus, and ii) to detect a 1.3 Mb region without recombination around the mating-type locus, through analysis of differentiation between the mating-type chromosomes. We observed three evolutionary strata, with significantly different levels of differentiation between mating-type chromosomes, suggesting a stepwise extension of recombination suppression. We detected no increase in transposable element content, non-synonymous substitutions or rearrangements in the non-recombining region. Recombination suppression was confirmed by segregation analyses. Our findings indicate that the convergent evolution of an extremely high frequency of self-fertile dikaryotic spores together with recombination suppression around the mating-type locus has occurred in multiple pseudo-homothallic fungal species.

## RESULTS

### *High frequency of self-fertile spores in* Schizothecium tetrasporum

The study of a single strain of *Schizothecium tetrasporum* (also known as *P. tetraspora*; Figure 1) suggested that this species was pseudo-homothallic and produced self-fertile heterokaryotic ascospores similarly to *P. anserina* (21,28), *i*.*e*., with most asci containing four dikaryotic spores, probably carrying both mating types. We studied the breeding system of *S. tetrasporum* further, by testing different media and identifying those optimal for reproducible and fast growth, and those optimal for sexual reproduction (see Material & Method section). Raju & Perkins found that the FGSC7436 strain produced 2% five-spore asci. In this study, we found that 10% of the asci produced by the CBS815.71 strain had five spores. We also observed, in rare frequencies, other kinds of asci, including six-spore asci, and some with “extra-large” ascospores, probably encompassing three nuclei, rather than the two usually encompassed during ascospore delimitation (Figure 1). The resulting mycelium produced numerous asexual single-cell propagules, for which we were unable to find appropriate conditions for germination and development. These cells probably, therefore, corresponded to spermatia rather than conidia, *i*.*e*. they are sexual cells acting as fertilizing elements and are unable to germinate, in contrast to asexual spores. *Podospora anserina* also produces spermatia, which function exclusively in the fertilization of the female structures, the protoperithecia (33). We generated an F1 progeny from the CBS815.71 dikaryotic strain, and isolated two homokaryotic offspring of opposite mating types, referred to hereafter as CBS815.71-sp3 and CBS815.71-sp6. The mating types of these strains were named according to the standard nomenclature (34): *MAT1-1* for CBS815.71-sp6 and *MAT1-2* for CBS815.71-sp3.

**Figure 1:**
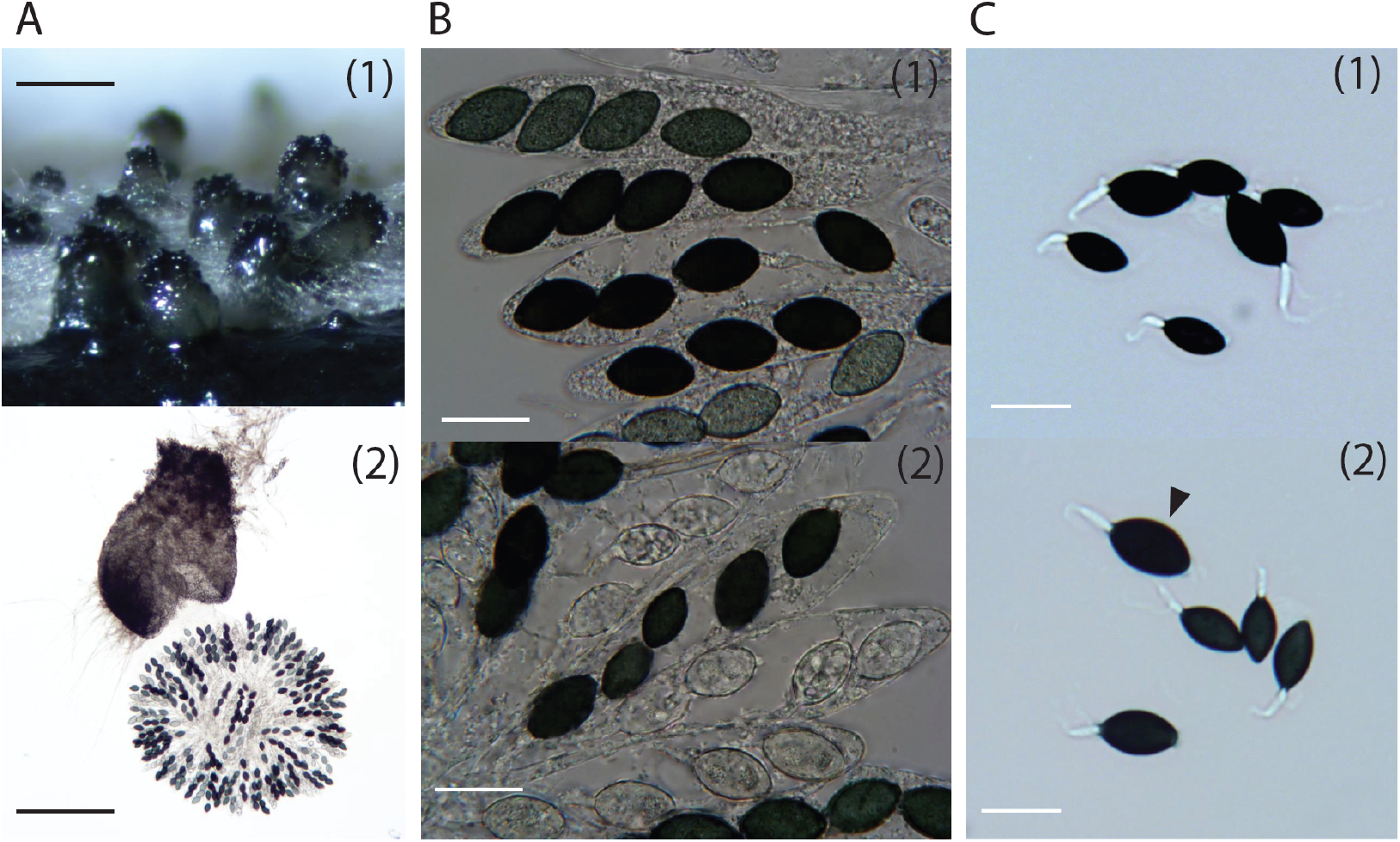
Sexual reproductive structures and ascospores of *Schizothecium tetrasporum* CBS815.71. **A**. Perithecia on *Miscanthus* (1) and squashed perithecia showing the rosette of asci (2); bars = 200 µm. **B**. Enlargements of asci within the rosettes (1) and a five-spore ascus with three dikaryotic ascospores and two monokaryotic ascospores (2); bars= 20 µm. **C**. Other abnormal asci observed in CBS815.71: (1) a six-spore ascus with four monokaryotic ascospores (small) and two dikaryotic ones (large); (2) a five-spore ascus with three monokaryotic ascospores (the three small spores in the middle), one dikaryotic (the large ascospore at the bottom) and another, probably trikaryotic, ascospore (arrowhead showing the very large ascospore); bars= 20 µm.

We assessed the frequency of second-division segregation (SDS) at the mating-type locus (Figure 2), by analyzing the progeny of a cross between CBS815.71-sp3 and CBS815.71-sp6. We collected 190 four-spore asci (each ascus containing the products of a given meiosis) and isolated large (dikaryotic) ascospores. Mating types were determined in the mycelium generated from germinating ascospores, by PCR with two primer pairs, each specific to one mating-type allele and yielding amplicons of different sizes. We initially genotyped only one ascospore per ascus, as this is usually sufficient to differentiate between two extreme cases: first-division segregation at meiosis (FDS) and second-division segregation (SDS). FDS yields four-spore asci containing two ascospores of a single mating type and another two ascospores of the opposite mating type, whereas SDS results in four ascospores, all containing both mating types (Figure 2A). We found that 87% (166/190) of the thalli resulting from ascospore germination carried both mating types, indicating that they resulted from SDS, and 13% (24/190) carried a single mating type, and were therefore either haploid or had two nuclei of the same mating type. Indeed, thalli carrying a single mating type may result not only from FDS, but also from the loss of a nucleus during growth in thalli resulting from SDS (Figure 2B). We differentiated between these two situations by determining the alleles at the mating-type locus in thalli resulting from the germination of the three remaining spores from the 24 four-spore asci in which we found a single mating type in the first genotyped ascospore. Only one of the 24 asci analyzed proved to be a true FDS ascus (*i*.*e*., with two *MAT1-1* and two *MAT1-2* ascospores). The other 23 asci contained at least one ascospore carrying both mating types, indicating that the homokaryotic thalli growing from their other ascospores resulted from the loss of nuclei during growth after SDS in asci (Figure 2B). We estimated the overall frequency of SDS for mating type at 99.5% (189/190) in the CBS815.71 strain.

**Figure 2:**
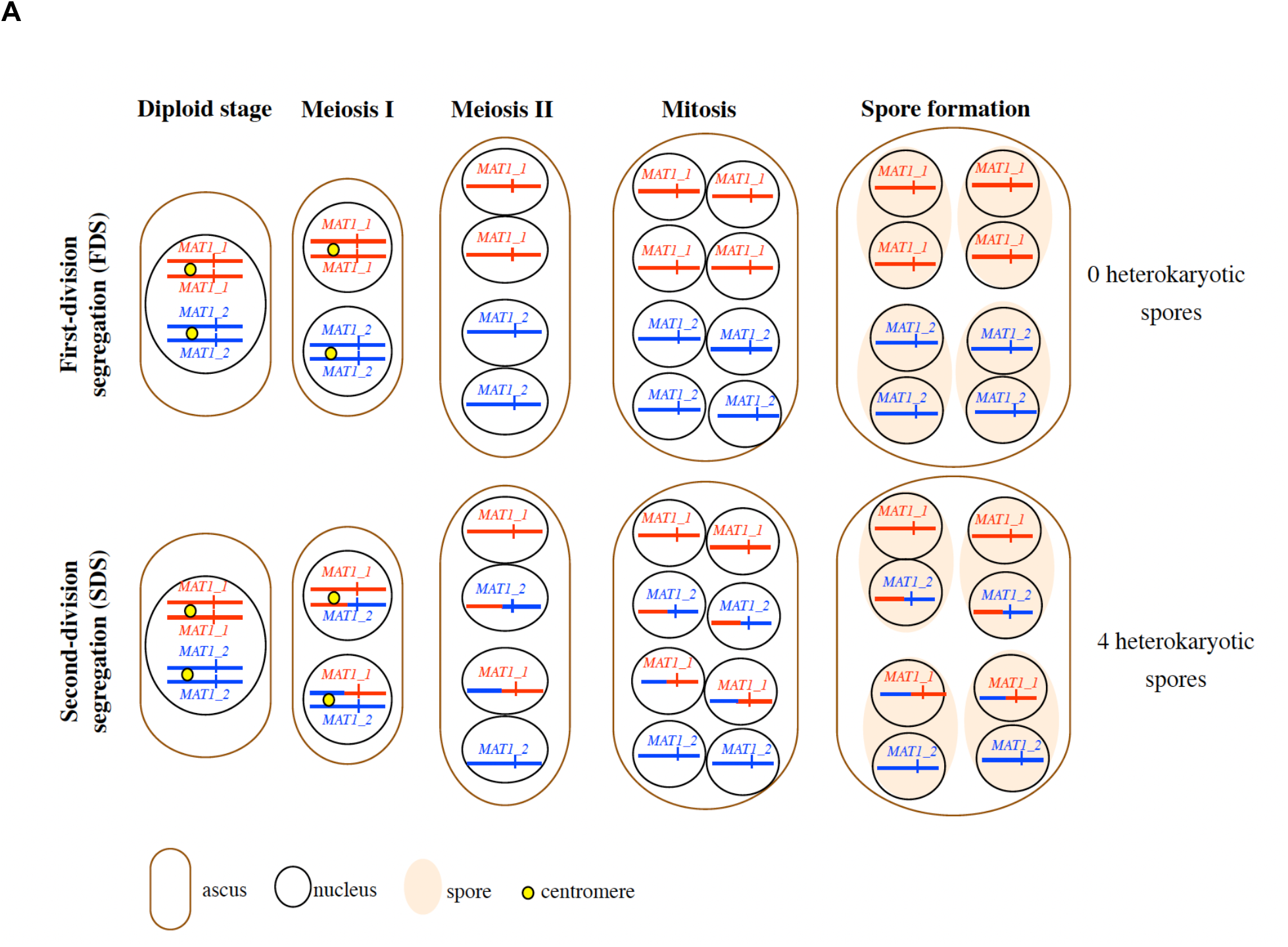
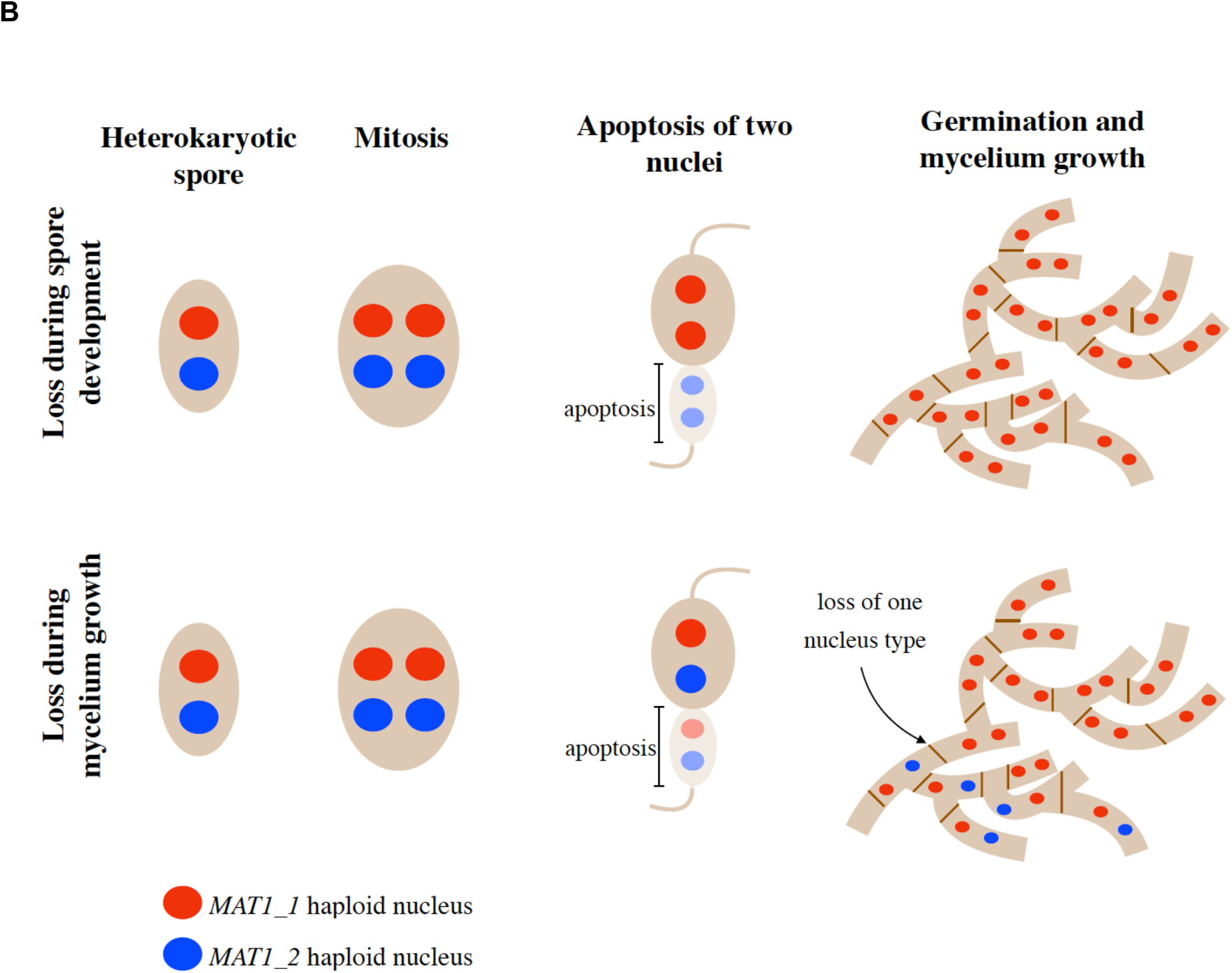
Segregation of the mating-type locus during meiosis, ascus composition and hypothetical scenarios for the loss of one mating type in thalli of *Schizothecium tetrasporum*. **A**. Theoretical composition of an ascus according to meiotic segregation pattern at the mating-type locus: if alleles at the mating-type locus (represented as a red or blue vertical bar) segregate during the first meiotic division (FDS, top), two *MAT-1-1* spores and two *MAT-1-2* spores are formed; conversely, if alleles at the mating-type locus segregate during the second meiotic division (SDS, bottom), four heterokaryotic spores carrying both mating types are formed. Only the chromosomes carrying the mating-type locus are represented, with their centromeres. The *MAT-1-1* and *MAT1-2* alleles at the mating-type locus are represented by red and blue vertical bars, respectively. Centromeres are represented by yellow circles at the diploid and meiosis I stages. Chromosomal arms are colored in red or blue according to the parental origin of the chromosomal fragment. **B**. Two hypothetical scenarios for the loss of one mating type despite SDS at the mating-type locus. SDS at the mating-type locus generates a spore carrying two nuclei of opposite mating types, but one nucleus may be lost during spore development (top) when half the nuclei gather in a region of the spore that subsequently undergoes apoptosis, or during mycelium growth (bottom).

We detected a substantial number of homokaryotic thalli originating from SDS asci (23 thalli from 189 germinating heterokaryotic ascospores), suggesting that the loss of nuclei corresponding to one mating type in heterokaryotic thalli may occur at a non-negligible frequency in the CBS815.71 strain. Raju and Perkins (21) showed that the two nuclei present in developing ascospores undergo mitosis, which is rapidly followed by the migration of two of the four daughter nuclei into the future pedicel cell, which are presumed to abort, or at least to fail to contribute to the nuclear composition of the resulting culture. It is therefore possible that homokaryotic thalli result from the migration of two nuclei carrying the same mating type into the future pedicel (leaving two nuclei of the same mating type in the maturing large ascospore cell), rather than the migration of two nuclei carrying opposite mating types (leaving two nuclei of opposite mating types in the maturing large ascospore cell; Figure 2B). It is difficult to test this hypothesis, because there is no easy way to follow the migration of nuclei during ascospore maturation. Alternatively, *MAT1-1/MAT1-2* heterokaryons may sometimes lose one type of nucleus during vegetative growth.

We tested this second hypothesis, by determining mating type in several sections of heterokaryotic thalli. A heterokaryon was generated by vegetative fusion (*i*.*e*., by mixing identical amounts of ground mycelia of the two homokaryotic strains of opposite mating types, CBS815.71-sp3 and CBS815-sp6). This procedure of mixed homokaryons chopped in very fine grain leads to the generation of a heterokaryotic mycelium through multiple vegetative fusions (35) and nuclear movements in the mycelial interconnected network of cells (36); no sexual reproduction can occur under these growth conditions. The resulting heterokaryotic mycelium was grown on V8 medium and incubated until the colony had reached 4 cm in diameter. It was then sliced into 70 pieces, each with a volume of about 1 mm^3^. PCR analyses demonstrated that only 31 of these 70 pieces carried both mating types, 29 contained only *MAT1-2* and 10 contained only *MAT1-1*. In a parallel experiment on another thallus of the same heterokaryon, the 70 pieces of mycelium were used to inoculate fresh M2 medium, and regenerating thalli were allowed to grow to a diameter of 1 cm. A randomly selected 1 mm^3^ piece of each of these thalli was sampled at the edge of growing colonies and genotyped by PCR for mating type. We found that 34 of the 70 regenerated thalli carried both mating types, whereas 25 carried only *MAT1-2* and 11 carried only *MAT1-1*. These results indicate that one of the two nuclei can be lost during vegetative growth, and they confirm the high rate of SDS in *S. tetrasporum*, yielding 99.5% of asci with all four ascospores initially containing both mating types, but with the occasional loss of one nucleus during vegetative growth. The *MAT1-1* mating type was preferentially lost in both experimental set-ups (chi^2^=9.25, *p*<0.005 for the first experiment and chi^2^=4.44, *p*<0.025 in the second experiment).

We investigated whether this loss of nuclei could be due to the relative fitness of the two types of nuclei, by investigating whether the two CBS815.71 homokaryons, carrying either the *MAT1-1* or *MAT1-2* mating type, displayed differences in growth rate or biomass production, and comparing them with the *MAT1-1/MAT1-2* heterokaryon. We measured growth rates on three different media and determined the dry weight obtained after growth on M2 medium for both homokaryons and the heterokaryon (Table S1A). No significant differences in growth or biomass were found between the homokaryons or between the homokaryons and the heterokaryon, the only significant effect being that of culture medium on growth rate (ANOVA, Table S1B). The growth rate of the *MAT1-1* homokaryon, corresponding to the mating type most often lost during growth, was slightly lower, although not significantly so (Table S1A).

### *Reference genome assembly for* Schizothecium tetrasporum

Using both Illumina and Oxford Nanopore MinION technology, we sequenced the genomes of the two homokaryotic strains of opposite mating types, CBS815.71-sp3 and CBS815.71-sp6. We built a *de novo* long read-based genome assembly for each, the first such assemblies to have been generated for *S. tetrasporum*, to our knowledge. For CBS815.71-sp3, we also had an Illumina-based assembly available that had been obtained before the long-read assemblies. BLAST searches and analyses of Illumina read coverage against the CBS815.71-sp3 long read-based assembly suggested the presence of bacterial contamination in the Nanopore data, but not in the Illumina data. We discarded scaffolds with high similarity scores to bacterial genomes in BLAST analyses for further analyses. The resulting long read-based genome assemblies for CBS815.71-sp3 and CBS815.71-sp6 contained 16 and 23 contigs and were 32.88 Mb and 32.83 Mb long, respectively. We found 96% of the 3,817 highly conserved orthologs in Sordariomycota from BUSCO analyses on the assemblies (Table S2). In the CBS815.71-sp3 assembly, seven contigs were more than 3.5 Mb long and contained all the conserved BUSCO orthologs. One contig corresponded to the mitochondrial genome and the other 8 contigs were less than 70 kb long, which is consistent with the expected number of chromosomes in Sordariales, an order in which several species have seven chromosomes (28,37). Telomeric repeats were present at both ends of contigs 1, 2, 4 and 5 in the CBS815.71-sp3 assembly, indicating that these contigs represent four entirely assembled chromosomes (Figure 3A). For CBS815.71-sp3, we compared the long read-based assembly obtained with the more fragmented Illumina-based assembly and found high levels of similarity and synteny (Figure S1). We used the long read-based assembly of CBS815.71-sp3 for further analyses. We found a high overall level of collinearity between the two homokaryotic long read-based genome assemblies for CBS815.71-sp3 and CBS815.71-sp6 (Figure 3B). The 2305 indels between assemblies that we detected were scattered throughout the genome and all were less than 10 bp long. These small indels were probably due to an artifactual enrichment of low-complexity regions in homopolymers in the Nanopore data and were not considered for subsequent analyses.

**Figure 3:**
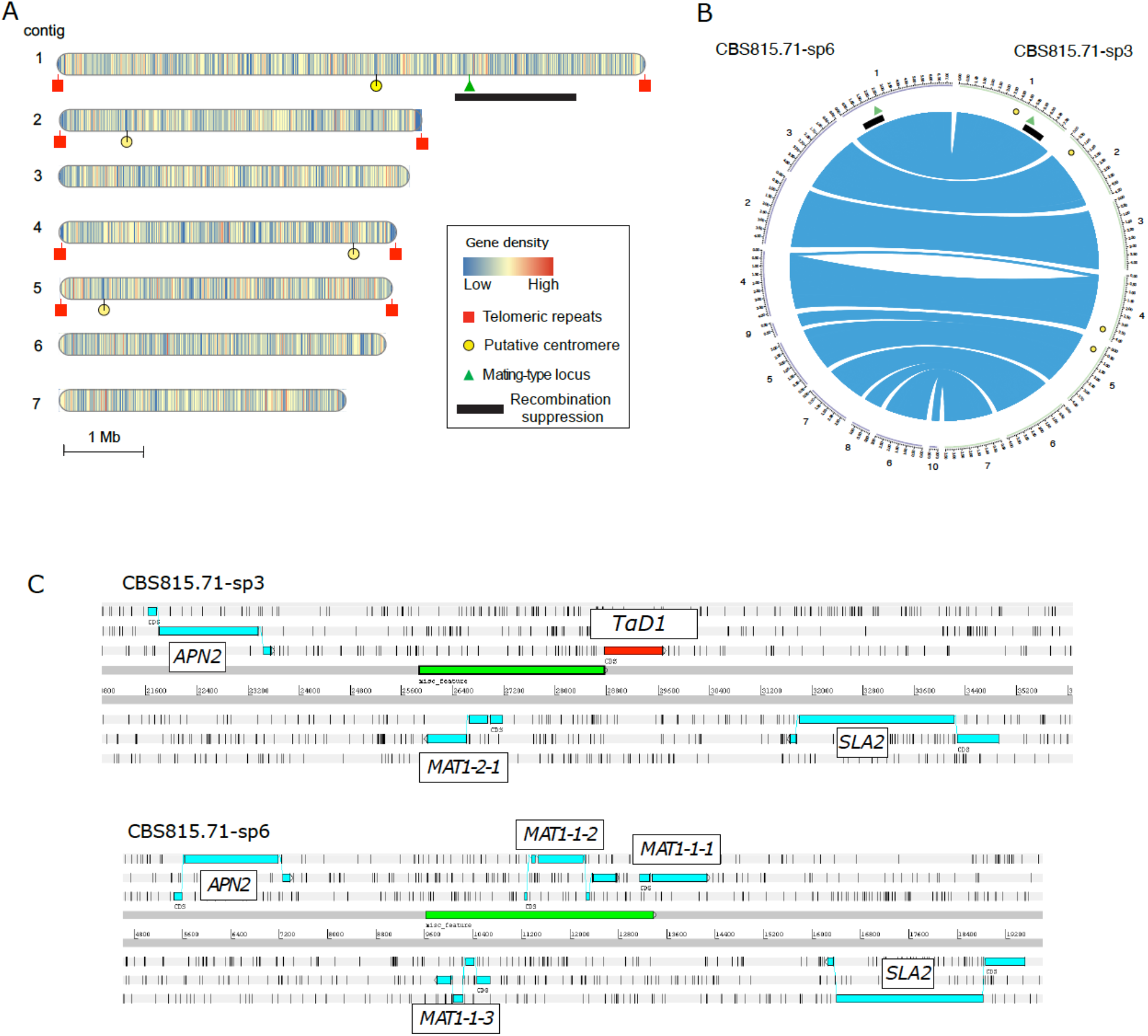
Genome features and annotation of the mating-type locus in the *Schizothecium tetrasporum* CBS815.71 strain. **A**. Ideograms representing the seven largest contigs of the CBS815.71-sp3 genome assembly. The mating-type locus is indicated with a green triangle, telomeric repeats with red boxes and putative centromeres with yellow circles. Gene density in 10 kb non-overlapping windows is represented by vertical bars on the contigs and the heatmap colors indicate low to high density. The non-recombining region around the mating-type locus is indicated by a black rectangle. **B**. Genome collinearity between opposite mating types of the *Schizothecium tetrasporum* CBS815.71 strain. Circular plot showing contigs from the CBS815.71-sp3 genome (green, right) and the CBS815.71-sp6 genome (purple, left). Only the largest contigs (>500 kb) were plotted. Blue links indicate the locations of orthologous genes (Mb). The mating-type locus is indicated with a green triangle. Putative centromeres and the non-recombining region around the mating-type locus identified on the CBS815.71-sp3 genome assembly are indicated by yellow circles and a black rectangle, respectively. **C**. Organization of the mating-type locus in CBS815.71-sp3 (top) and CBS815.71-sp6 (bottom) assemblies viewed in Artemis v18.0.2 (100). The *APN2* and *SLA2* genes flanking the mating-type locus are indicated. The green boxes represent the nucleoide sequences differing between the two idiomorphs that were used to design markers for assessing mating type in the progeny. The *MAT1-1-1* pseudogene (*TαD1*) present in the CBS815.71-sp3 (*MAT1-2*) genome is boxed in red. The three forward and three reverse frames of the DNA sequence are presented; vertical bars indicate putative STOP codons.

Repetitive sequences accounted for about 7% of the assemblies. In the CBS815.71-sp3 assembly, we identified 182 transposable element (TE) family consensus sequences: 4.2% Gypsy LTRs and 6.7% other LTRs, 7% non-LTR SINEs and LINEs, 7.5% TcMar-Fot1 DNA transposons, 3.4% other DNA transposons and the remaining 70.5% repetitive sequences could not be annotated (Figure S2A). Both genomes contained very few (<0.1 %) signatures of RIP (repeat-induced point mutations), a fungal genome defense against transposable elements inducing C to T mutations in repeated sequences at target dinucleotides (38,39).

### *Structure of the mating-type locus in* Schizothecium tetrasporum

The mating-type locus was located at about the 5.180 Mbp position on contig 1 of the CBS815.71-sp3 assembly, which was considered likely to correspond to a fully assembled chromosome (Figure 3). The mating-type locus of *S. tetrasporum* had the typical structure of heterothallic Sordariales (20), with two highly divergent “alleles”, referred to as idiomorphs, flanked by the *APN2* and *SLA2* genes (Figure 3C; Figure S3, (40,41)). The *MAT1-2* idiomorph, present in CBS815.71-sp6, harbored a single functional gene (*MAT1-2-1*), orthologous to the *mat+ FPR1* (*Pa_1_20590*) gene of *P. anserina* and the *mat_a-1* gene of *Neurospora crassa*. The other idiomorph, *MAT1-1*, carried three genes (*MAT1-1-1, MAT1-1-2* and *MAT1-1-3*), orthologous and syntenic with the three genes present in the *mat-* idiomorph of *P. anserina* (*Pa_1_20593, _20592, _20591*) and the *mat_A* idiomorph of *N. crassa (NCU01958, NCU01959, NCU01960). MAT1-1-1* contained the typical MATα_HMG domain (pfam04769) and *MAT1-1-2* contained a HPG domain (pfam17043). *MAT1-1-3* and *MAT1-2-1* carried a MATA_HMG domain (cd01389). The 3’ end of the *MAT1-1-1* coding sequence was found to extend outside the *MAT1-1* idiomorph in *S. tetrasporum*, as shown by its presence next to the *MAT1-2* idiomorph. This truncated *MAT1-1-1* gene close to the *MAT1-2* idiomorph has a putative open reading frame starting 68 residues downstreaming of the start codon of the typical *MAT1-1-1* gene structure (Figure S3). However, this putative start codon is in a poor context for translation initiation, with a pyrimidine in position −3 (40). The truncated *MAT1-1-1* gene encodes most of the MATα_HMG domain, and the second conserved domain adjacent to the MATα_HMG domain (Figure S3; (42)). This gene was therefore named *TαD1* (truncated MAT<u>α</u>_HMG <u>d</u>omain). This name does not follow the standard mating-type gene nomenclature (34), as it is not included in the mating-type locus. TαD1 may be a pseudogene and its transcriptional status is unknown in *S. tetrasporum*. A similar truncated *MAT1-1-1* gene is present and has little or no expression in *MAT1-2* strains in the Chaetomiaceae (Sordariales) *Myceliophthora heterothallica, M. fergusii, M. hinnulea* and *Humicola hyalothermophila* (43). It is unknown whether *TαD1* has any role in sexual reproduction.

### A region with stepwise recombination suppression around the mating-type locus in Schizothecium tetrasporum, located 1 Mb from the centromere

We plotted the synonymous divergence (dS) between the two genomes of opposite mating types for each gene along the contigs of the CBS815.71-sp3 assembly (Figure 4; Figure S4). Most of the dS values obtained were zero, as expected in a selfing species with a high degree of homozygosity. Only around the mating-type locus did we find a cluster of genes with high dS values (Figure 4), indicating recombination suppression. The high degree of homozygosity throughout the genome indicates that many selfing events occurred in the ancestors of the CBS815.71 heterokaryotic strain; only the regions fully linked to the mating-type locus have remained heterozygous under selfing (27). Indeed, for any gene linked to the mating-type locus, alleles associated with the alternative mating types gradually accumulate different mutations over the time since recombination suppression. Synonymous divergence (dS) can, thus, be used as a proxy for the age of the linkage between a gene and the mating-type locus (7).

**Figure 4:**
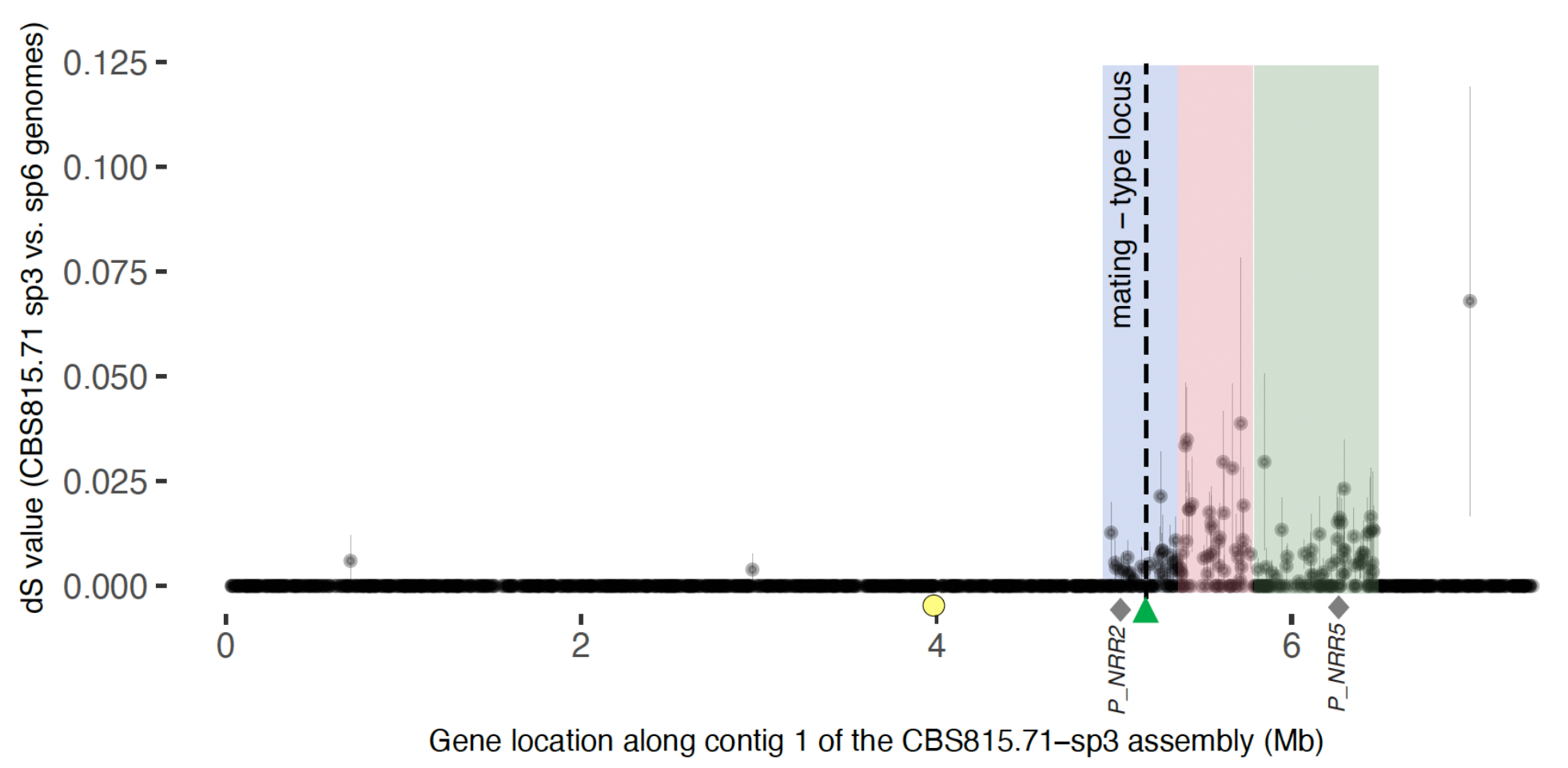
Synonymous divergence (dS) along the mating-type chromosome in the *Schizothecium tetrasporum* CBS815.71 strain. dS was calculated per gene, between the orthologs common to the CBS815.71-sp3 and CBS815.71-sp6 assemblies and was plotted, according to the CBS815.71-sp3 assembly gene position (contig 1), along the *x-*axis. Evolutionary strata 1, 2, and 3 are indicated by red, green and blue rectangles, respectively. The mating-type locus is indicated with a black dotted vertical line and a green triangle. The putative centromere is indicated with a yellow circle. The *P_NRR2* and *P_NRR5* markers, which displayed no recombination event with the mating-type locus, are indicated with gray diamonds.

The cluster of genes with high dS values around the mating-type locus spanned a 1.47 Mb region located between 4.983 Mbp and6.457 Mbp on scaffold 1 of the CBS815.71-sp3 assembly, encompassing 406 genes in the CBS815.71-sp3 assembly and 413 genes in the CBS815.71-sp6 assembly. The dS values were lowest in the center of the non-recombining region (Figure 4), suggesting the occurrence of gene conversion. We found no predicted function potentially directly related to mating compatibility for any of the 406 genes in the non-recombining region of the CBS815.71-sp3 assembly other than the mating-type locus. A total of 15 genes in the non-recombining region encoded proteins that were identified as putative transcription factors or involved in signaling pathways (Table S3). The four orthologs of the *P. anserina* genes located in its non-recombining region and known to be involved in post-mating sexual development (but not mating compatibility) were also found in the non-recombining region in *S. tetrasporum*: *ami1*, controlling nuclear migration events (44), *PaRID*, a DNA methyltransferase controlling sexual development in *P. anserina* (45), *AS4*, encoding the eEF1A translation elongation factor and *su12*, encoding a ribosomal protein, mutations of which often result in sterility in *P. anserina* (46,47).

The d_S_ values plotted along the mating-type chromosome seemed to indicate the existence of three distinct evolutionary strata (Figure 4; Figure S6). We tested the existence of evolutionary strata more formally, by splitting the heterozygous region with sliding limits in the CBS815.71-sp3 assembly and calculating, for each limit, the difference in dS values between the two segments; limits between strata should correspond to a peak in dS value differences according to stratum definition (27). We first arbitrarily set the limit of the first stratum by eye at 5,350,000 bp and applied the automatic procedure to the rest of the heterozygous region. The dS values peaked at 5,768,335 bp, identifying the limit between strata 1 and 2 (Figure S6). We then re-applied the procedure to the part of the heterozygous region without stratum 2. The dS values peaked at 5,388,708 bp, identifying the limit between strata 1 and 3 (Figure S6). The median dS values were significantly different between strata 1 and 2 (pairwise Wilcoxon test *p*-value = 0.00134 with the False Discovery Rate (FDR) correction) and between strata 1 and 3 (pairwise Wilcoxon test *p*-value = 0.00015 with FDR correction), supporting the existence of distinct evolutionary strata. Median dS values did not differ significantly between strata 2 and 3, located on opposite sides of stratum 1 (pairwise multiple-comparison Wilcoxon test *p*-value = 0.27571 with FDR correction). Strata 1 and 3 each spanned 400 kb, and stratum 2 spanned about 700 kb. The most ancient evolutionary stratum, stratum 1, with the highest dS values, does not encompass the mating-type locus. The mating-type locus was located in the middle of stratum 3 (Figure S6).

We looked for the centromere on the mating-type chromosome because its position may provide important insight into the segregation of the mating-type locus at the first meiotic division. We found a single location with a drop in GC content along the mating-type chromosome of the CBS815.71-sp3 assembly outside of the telomeric regions, with 53.5% GC between 3.990 Mbp and 4.025 Mbp, versus the genome-wide mean value of 58.76% (Figure S5). A drop in GC content can pinpoint the centromere in some fungi, such as some *Neurospora spp*. in which GC content drops to about 44%, whereas the genome-wide mean value is about 53% (48). The lower variation of GC content observed in the CBS815.71-sp3 genome assembly may be due to a lack of RIP in *S. tetrasporum*. For the other three fully assembled chromosomes of the CBS815.71-sp3 genome, similar drops in GC content were also identified at a single location per chromosome, co-localizing with clusters of repeats. We characterized the repeat content of these four putative centromere regions. We found no specific repeats, but we did detect an enrichment in *TcMar-Fot1* DNA transposons relative to the genome-wide mean (Figure S2A). The putative centromere of the mating-type chromosome was located about 1 Mb from the edge of the non-recombining region around the mating-type locus.

We found no enrichment in repetitive sequences in the non-recombining region around the mating-type locus relative to autosomes and the recombining regions on the mating-type chromosome in the CBS815.71-sp3 assembly. The mean repetitive sequence densities calculated in 10 kb windows overlapping by 1 kb were 2.52%, 5.05% and 4.21% in the non-recombining region around the mating-type locus, the autosomes and the recombining regions on the mating-type chromosome, respectively; pairwise Wilcoxon tests revealed no significant differences in repetitive sequence densities. We found no difference in TE family proportions between these regions either (Figure S2B). Repetitive sequence density was significantly higher in stratum 2 than in stratum 3 (*p*-value for a pairwise Wilcoxon test = 0.0053 with FDR correction). We found no genes with a dN/dS ratio >1 in the non-recombining region, i.e. with higher non-synonymous than synonymous substitution rates (the mean dN/dS was 0.11 for genes with positive dS values). As noted above, we found no evidence for genomic rearrangements between the two mating-type chromosomes in the non-recombining region (Figure 3B).

### Validation of the region without recombination by progeny analysis

We investigated the lack of recombination in the heterozygous region around the mating-type locus in the CBS815.71 strain. We obtained offspring resulting from a selfing event in the CBS815.71 strain, *i*.*e*., a cross between CBS815.71-sp3 and CBS815.71-sp6. We genotyped 107 offspring resulting from haploid spores isolated from five-spore asci (i.e., asci with three large dikaryotic spores and two small monokaryotic spores; Figure 1). We initially used two genetic markers, one at each end of the heterozygous region, and then three additional markers in the three recombinant offspring to delimit more finely the regions in which recombination events occurred (Figure S7). We detected no recombination events at the edge of stratum 3 (*i*.*e*., between the *P_NRR2* marker and the mating-type locus) but we found recombination events at the edge of stratum 2 (between the markers *P_NRR4* and *P_NRR5*) in three of the 107 offspring. We found no recombination event between the mating-type locus and the additional markers *P_NRR7, P_NRR6* and *P_NRR5* located 0.808 Mb, 1.054 Mb and 1.088 Mb from the mating-type locus, respectively. Given the position of the MAT locus, this suggests a complete lack of recombination over a 1.3 Mb region in strata 1 and 3, and in a large part of stratum 2, as recombination occurred only at the edge of this stratum. Stratum 2 is, therefore, likely slightly smaller than the heterozygous region in the CBS815.71 strain, the heterozygosity at its edge probably resulting from an outcrossing event a few generations ago.

### Orthologs in the non-recombining regions across Sordariales

We aimed at investigating gene orthology relationships and synteny between the non-recombining regions on the mating-type chromosomes of the three pseudo-homothallic species, *S. tetrasporum, P. anserina* and *N. tetrasperma*. However, several rearrangements occurred following recombination suppression on the mating-type chromosomes in *N. tetrasperma*. We therefore used the genome of the closely related heterothallic *N. crassa* species instead, as a proxy for the ancestral gene order in *N. tetrasperma* before recombination suppression. No recombination suppression around the mating-type locus has been reported in *N. crassa* (26,49). This analysis therefore also provided an opportunity for an interesting contrast with species displaying recombination suppression. In an analysis of single-copy orthologs, we found that the mating-type chromosomes, and all the other chromosomes, displayed regions of synteny, but also breaks of synteny between species, and mesosynteny, *i*.*e*., a global conservation of gene content despite rearrangements in order and orientation (Figure 5A; Figure S8), as expected given their phylogenetic distance (50). In particular, 69 orthologs common to the species studied and belonging to stratum 3 of *S. tetrasporum* had a well-conserved gene order in a 300 kb region around the mating-type locus (Figure 5A), suggesting some conservation of the ancestral gene order in the region around the mating-type locus in these distantly related pseudo-homothallic species, and in other non-pseudo-homothallic Sordariales, as previously reported (32). The region around the putative centromere identified on the *S. tetrasporum* mating-type chromosome had orthologous genes and gene orders conserved with the *P. anserina* and *N. crassa* centromeric region over a distance of 700 kb despite an inversion, providing support for the putative location of the centromere on the *S. tetrasporum* mating-type chromosome identified above (Figure 5A).

**Figure 5:**
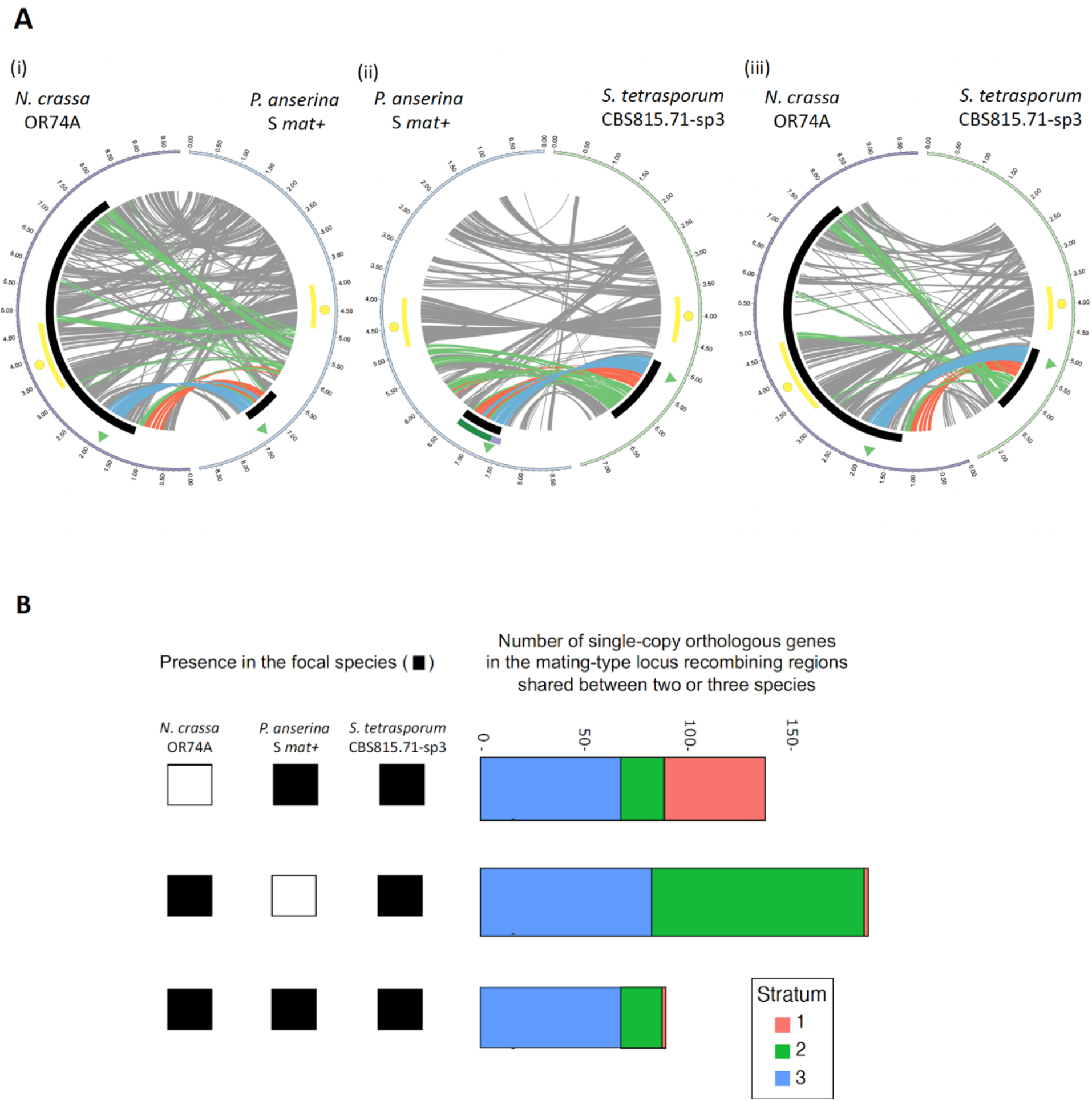
Collinearity and shared gene content between the mating-type chromosomes of *Schizothecium tetrasporum, Podospora anserina* and *Neurospora tetrasperma*. **A**. Collinearity between (i) *N. crassa* (left) and *P. anserina* (right); (ii) *P. anserina* (left) and *S. tetrasporum* (right); (iii) *N. crassa* (left) and *S. tetrasporum* (right). Collinearity is based on single-copy orthologous genes. Centromere is indicated with a yellow circle and the region of well-conserved gene order around the centromere is highlighted with a yellow rectangle. The mating-type locus location is shown with a green triangle and the non-recombining region around the mating-type locus in *P. anserina* (23) and *S. tetrasporum* with a black track. Along the recombining mating-type chromosome of *N. crassa* a black rectangle indicates the orthologous region that is non recombining in multiple *N. tetrasperma* lineages (26). Red, green and blue links highlight orthologous genes located in the evolutionary strata 1, 2, and 3 of *S. tetrasporum*, respectively. Grey links indicate other orthologous genes. On panel (ii), the evolutionary strata identified previously in the *P. pseudocomata* CBS415.72 strain, that is closely related to the *P. anserina* S strain, are indicated by purple (for the oldest stratum) and dark green (for the youngest stratum) tracks (27), at left. B. Number of single-copy orthologous genes present in the non-recombining region of *S. tetrasporum* that are also in the non-recombining regions of *P. anserina* and/or *N. tetrasperma*, as indicated by the squares at the bottom (a filled square indicates the presence in the non-recombining region of the focal species).

We also found conserved gene content and synteny between parts of the non-recombining regions in the three species despite rearrangements. In *S. tetrasporum*, 261 of the 406 genes located in the identified region of suppressed recombination around the mating-type locus had orthologous genes in both *P. anserina* and *N. crassa*. We found that 138 of these 261 genes were also located in the region of suppressed recombination around the mating-type locus in the *P. anserina* S strain (Figure 5B). Most of these shared genes (85%) were part of stratum 3, encompassing the mating-type locus, and stratum 1, the oldest evolutionary stratum in *S. tetrasporum* (Figure 5A(2),5B). No evolutionary strata have previously been identified in *P. anserina*, but the oldest evolutionary stratum in *P. pseudocomata*, a cryptic species closely related to *P. anserina* (27), was orthologous to stratum 3 in *S. tetrasporum*. Most genes (82%) of stratum 2 in *S. tetrasporum* had orthologs in *P. anserina* in the recombining region between the mating-type locus and the centromere (Figure 5A(ii) & 5B). The region of recombination suppression was located from the mating-type locus in the direction of the telomere in *S. tetrasporum*, whereas it was located from the mating-type locus in the direction of the centromere in *P. anserina* and other species of the *P. anserina* species complex (Figure 5A(ii)&5B). The distance between the centromere and the mating-type locus was 1.1 Mb in *S. tetrasporum* and 2.9 Mb in *P. anserina*.

We further found that 188 of the 406 genes in the non-recombining region of *S. tetrasporum*, (mostly in strata 3 and 2) were also present in the non-recombining region in *N. tetrasperma* (Figure 5B), with the genes from *S. tetrasporum* stratum 1 present in the flanking region of the non-recombining region in *N. tetrasperma*. In total, 90 genes were common to the non-recombining regions of the three pseudo-homothallic species, including genes at the two edges of these non-recombining regions (Figure 5A). The predicted functions of these 90 orthologs, when available, did not appear to be related to mating or mating-type functions (Table S4).

## DISCUSSION

### Striking evolutionary convergence

This study of the breeding system and sexual biology of the little-studied pseudo-homothallic fungus *S. tetrasporum* reveals a striking convergence with *P. anserina* and other cryptic species of the *P. anserina* complex, for multiple ecological and reproductive traits. These two distantly related fungi thrive in herbivore dung and produce predominantly dikaryotic spores carrying opposite mating types. In both cases, this is due to mating-type segregation during the second meiotic division associated with a single systematic CO event between the centromere and the mating-type locus. In addition, both *S. tetrasporum* and *P. anserina* display recombination suppression around this locus ((23) and this study). We found that recombination suppression had expanded in several successive steps in *S. tetrasporum*, as in *P. peudocomata*, which belongs to the *P. anserina* species complex (27). The association between pseudo-homothallism (self-fertile dikaryotic spores), recombination suppression around the mating-type locus and evolutionary strata also evolved independently in the *N. tetrasperma* complex, representing an additional case of convergence. However, unlike *S. tetrasporum* and *P. anserina, N. tetrasperma* displays recombination suppression up to the centromere and mating-type segregation during the first meiotic division (26,48).

### Association between pseudo-homothallism and recombination suppression: evolutionary and proximal causes

Such a convergent association between recombination suppression and pseudo-homothallism (with the production of heterokaryotic spores carrying both mating types and growth as a heterokaryotic mycelium rather than as a homokaryon in the predominant phase of the lifecycle, as in most ascomycetes) suggests causal relationships. Suppression of recombination around the mating-type locus may be selected for if it favors the production of heterokaryotic ascospores, thus facilitating the compaction of nuclei of opposite mating types in dispersing spores under the particular pattern of meiotic segregation. The resulting self-fertile heterokaryotic ascospores allow reproductive insurance in short-lived habitats such as dung. However, the occurrence of a CO between the centromere and the proximal end of the region of suppressed recombination in almost all meiosis I to cause mating-type segregation during the second meiotic division still calls for an explanation. Indeed, if we consider the 21 ± 3 COs occurring at each meiosis in one nucleus of *Sordaria macrospora* (51,52) as a proxy for the number of COs occurring in *S. tetrasporum* displaying approximately the same genome size, and if we take into account the number of obligate COs occurring on each of the seven chromosomes, we can estimate that 11 to 17 COs remain to be randomly distributed across a genome of 32 Mbp. It is therefore unlikely that one CO takes place between the centromere and the mating-type locus in 99.5 % of the meioses in *P. anserina*, the distance between the centromere and the mating-type locus being 2.9 Mbp, and in *S. tetrasporum*, the distance between the centromere and the mating-type locus being 1.2 Mbp. The occurrence of a second CO between the centromere and the mating-type locus should be prevented, as two COs in this region would result in FDS. This second CO might be prevented by positive interference, but this mechanism alone may not be sufficient to reach the very high frequency of SDS.

It therefore appears more likely that a yet unidentified mechanism has evolved that enforces the occurrence of the single and systematic CO event between the centromere and the mating-type locus, independently in both *P. anserina* and *S. tetrasporum*, together with a mechanism compacting two nuclei per ascospore, for the production of self-fertile ascospores. Suppression of recombination around the mating-type locus may be directly linked to the mechanism that prevents a second CO around the mating type locus, which would yield FSD instead of SDS. The selective advantage of producing self-fertile spores would then be the driver to develop this mechanism.

An alternative hypothesis may be proposed considering that the mating-type locus is always heterozygous in heterokaryons. Indeed, a recent theoretical model showed that recombination suppression capturing a permanently heterozygous locus may be selected due to its capacity to shelter deleterious alleles in organisms with an extended diploid or dikaryotic phase (14). More precisely, recombination suppression may be selected because it protects DNA fragments with fewer deleterious mutations than average in the population, but the load of these mutations will still prove problematic when they become frequent enough to become homozygous; the capture of a permanently heterozygous allele prevents this effect (14). This mechanism can also explain the evolution of successive evolutionary strata, as observed here in *S. tetrasporum* and, previously, in *N. tetrasperma* and *P. pseudocomata* (22,27,27,48), the basidiomycete mushroom *Agaricus bisporus* (53) and repeatedly in *Microbotryum* basidiomycete fungi, plant pathogens in natural ecosystems (8,9,18,54). This mechanism may also account for the suppression of recombination in *N. tetrasperma*.

In this study, we found no detectable differences in growth between homokaryons of the two mating types and the heterokaryon in *S. tetrasporum*, but we found that one of the mating-type alleles was more often lost during mycelial growth. In addition, fitness differences even smaller than those detectable here can promote the evolution of recombination suppression (14), and genes with deleterious alleles may be involved in specific parts of the life cycle not evaluated *in vitro*, and related to the dung habitat or resistance to harsh conditions. We found that 13% of the thalli grown from ascospores *in vitro* ended up homokaryotic, and the mechanism involved could function in haplodiplontic lifecycles, depending on the mean length or frequency of the haploid phase and the strength of selection (14). The mycelia in dung may not grow for long enough for the loss of a nucleus to be occur very often, especially as *S. tetrasporum* is mostly found on small pellets of rabbit dung (unpublished data).

The finding that the non-recombining regions in *P. anserina, N. tetrasperma* and *S. tetrasporum* contain a number of shared orthologous genes, particularly at the two edges of the non-recombining region, and that these regions display some level of synteny, raises the possibility of a third hypothesis: the selection of recombination suppression to maintain beneficial allelic combinations (55), as observed for supergenes in plants and animals (56,57). No function known to be directly related to mating compatibility was found among these orthologous genes, except for the mating-type locus, but several genes involved in sexual development in *P. anserina* are located in the non-recombining regions in both *P. anserina* (23) and *S. tetrasporum* (this study): *ami1*, controlling nuclear migration events (44) and *PaRID*, a RID-like putative cytosine methyltransferase that controls sexual development (45). However, there is no evidence of antagonistic selection between mating types for these functions (23) and we found no phenotypic differences between homokaryons of opposite mating types. Nevertheless, we cannot rule out the possibility that there are differences in other parts of the life cycle not investigated here *in vitro*.

These various evolutionary hypotheses for explaining recombination suppression are not mutually exclusive, and the evolutionary causes of recombination suppression may differ between evolutionary strata and species. The proximal (mechanistic) causes of recombination suppression also remain unknown. We detected no chromosomal inversion between mating-type chromosomes in *S. tetrasporum*, as in some other fungi in which recombination is also suppressed close to the mating-type locus without inversion, such as *P. anserina* and other species of the *P. anserina* complex, *N. tetrasperma* and *Microbotryum* fungi (8,23,26,27). Histone modification can also prevent recombination (58,59) and this aspect warrants investigation in *S. tetrasporum*. There may also be recombination modifiers in *cis*. The proximal cause for the single CO event between the centromere and the mating-type locus is also unknown. The shorter genomic region between the centromere and the mating-type locus in *S. tetrasporum* (1.2 Mb) than in *P. anserina* (2.9 Mb) makes this species an excellent model for identifying the proximal factors responsible for the single CO event, if they are located in *cis*. The convergent evolution of a single systematic CO event between the mating-type locus and the centromere in these two species provides an opportunity to study the genetic basis of the convergent evolution of this phenomenon.

### Evolutionary strata

We detected three evolutionary strata in the CBS815.71 *S. tetrasporum* strain, indicating a stepwise extension of recombination suppression. Our progeny analysis showed that stratum 2 did not cover the entire corresponding heterozygous region, the extremity of the heterozygous region probably being a remnant of an outcrossing event a few generations ago. Nevertheless, the progeny analysis showed that strata 1 and 3, and much of stratum 2, probably display a full suppression of recombination. This gradual progression of recombination suppression cannot be explained by sexual antagonism, as male and female functions are not associated with mating types in fungi (19,60). In addition, monokaryotic ascospores are rare in *S. tetrasporum* and traits and ecological features are similar between mating types, so other types of antagonistic selection are also unlikely. Stepwise recombination suppression in this dikaryotic fungus may have been selected to shelter deleterious alleles (14). It may appear surprising that the most ancient evolutionary stratum, stratum 1, with the highest dS values, does not encompass the mating-type locus. However, this oldest stratum is very close to the mating-type locus and the deleterious allele sheltering model predicts the possible evolution of recombination suppression at sites tightly linked to but not encompassing the permanently heterozygous locus (14). This situation has, in fact, been observed in the chestnut blight fungus *Cryphonectria parasitica* (61). It is also possible that the most ancient evolutionary stratum encompassed the mating-type locus but that the limits of the stratum were obscured by subsequent gene conversion events. Alternatively, the different strata may correspond to gradual increases in the SDS frequency, by stepwise increase of the efficiency of the mechanism preventing a second CO in the region of mating-type locus and FDS. The selective pressure for this stepwise increase would be to reach a higher number of dikaryotic ascospores growing into self-fertile mycelia.

### No sign of degeneration in the non-recombining region

We found no enrichment in repeats, rearrangements or non-synonymous substitutions, whereas transposable elements, non-synonymous substitutions and non-optimal codon usage have all been found to increase with time since recombination suppression in the mating-type chromosomes of *Microbotryum* anther-smut fungi (9,54,62). Gene disruptions due to small indels remain to be investigated. The suppression of recombination may have occurred too recently in *S. tetrasporum*, or recombination or gene conversion may occur occasionally, as suggested in *P. anserina* (27). The low dS values in the center of the non-recombining region, between strata 1 and 2, suggests that gene conversion may occur, as observed in the non-recombining regions of mating-type chromosomes in other fungi (8,9,23,63). We found no footprints of RIP, the fungal genome defense mechanism that introduces cytosine-to-thymine transitions into repetitive sequences to control transposable elements (38,39). Nevertheless, the TE load was not high in *S. tetrasporum*, suggesting that this fungus may have evolved another mechanism of transposable element control, such as RNA silencing (64).

### *Mating system in* Schizothecium tetrasporum

We found a higher percentage of homokaryotic thalli resulting from ascospore germination in *S. tetrasporum* than in *P. anserina* (13% versus 1%). Homokaryons may be recovered due to first-division segregation, the loss of one nucleus after germination or a particular pattern of nucleus migration following the post-meiosis mitosis. We found that nuclei could be lost during vegetative growth *in vitro* in *S. tetrasporum* (here in 13% of the germinating ascospores), whereas this is not the case in *P. anserina* (47), suggesting that *S. tetrasporum* may sometimes, albeit rarely, end up homokaryotic in nature. Such nucleus losses may be due to active competition between nuclei (65,66) or the presence of deleterious alleles associated with a mating type. The occurrence of mycelia of a single mating type could promote some degree of outcrossing, which could be checked in analyses of the levels of heterozygosity of dikaryons in natural populations. Nevertheless, the rate of self-fertile (heterokaryotic) ascospores was high, and this would be expected to promote a high level of selfing. Population genomics approaches detected widespread genome-wide homozygosity in dikaryons of *P. anserina*, indicating high selfing rates, but with occasional stretches of heterozygosity, also indicating the occurrence of outcrossing in nature (27,67,68). The optimal media and conditions for vegetative growth and sexual reproduction identified here will facilitate further investigations of the lifecycle and biology of *S. tetrasporum*. The reference strain was found to be highly homozygous, as expected under high rates of selfing. If we assume that the non-recombining region evolved to prevent a second CO, to maximize the frequency of SDS and thereby of self-fertile spores, it may appear intriguing that no mechanism evolved to subsequently maintain a dikaryotic mycelium, as for instance the crozier cells that are present in the perithecia and are analogous to the dikaryotic clamp cells of Basidiomycetes (69). It may be that, in nature, the mycelium does not grow long enough to frequently loose a nucleus or that the occurrence of mycelia carrying a single mating type is beneficial in promoting outcrossing, as does the formation of a few asci with 5 ascospores including two monokaryotic ascospores.

## Conclusion

In conclusion, we found a striking convergence in lifecycle, sexual biology and stepwise recombination suppression around the mating-type locus in phylogenetically distant fungi of the order Sordariales. Documenting convergent evolution is important, as it shows that evolution can be repeatable, and not only contingent (70). Studies of these traits in additional species would make it possible to perform a more robust assessment of the degree of convergence in trait association and, thus, of the causal relationship. Furthermore, the broad convergence towards an association between dikaryotic life cycle and stepwise recombination suppression beyond the mating-type locus despite the lack of sexual antagonism provides support for alternative hypotheses, such as deleterious allele sheltering, as an explanation for the suppression of recombination beyond mating-type or sex-determining alleles (14).

## MATERIALS AND METHODS

### Strain origins, experimental crosses and culture

The dikaryotic strain CBS815.71 of *S. tetrasporum* was obtained from the Centraalbureau voor Schimmelcultures (CBS). No optimized methods have been described for culturing *S. tetrasporum* and inducing sexual reproduction in this species. We therefore tested several media, to select those optimal for growth and sexual reproduction (M2, M3, Oat Meal Agar and V8 agar; (71)), and for ascospore germination (the same media plus *P. anserina* germination medium (G), (71)). Growth and sexual reproduction were optimal on V8 medium (composition per liter: 15 g agar, 2 g CaCO3, 200 mL *V8* tomato juice and 800 mL water) at 22°C with 12 h of illumination. Incubation at a higher temperature, such as 26°C, often resulted in aborted fruiting bodies. Growth and ascospore recovery were very slow at lower temperatures (*e*.*g*., 18°C). We observed 100% germination on G medium (composition per liter: 4.4 g ammonium acetate, 15 g bactopeptone and 13 g agar) with a 65°C heat shock for 30 minutes just after the plating of ascospores. On all the other media tested, germination rates were lower, with no germination in the absence of a heat shock. In optimal conditions, mature perithecia ejecting ascospores were obtained three weeks after inoculation on V8 medium, and germination occurred overnight on G medium after a heat shock. Ascospores could be collected and plated with the same methods as used for *P. anserina* (71).

In these optimal conditions, we generated and isolated two F1 monokaryotic strains by inducing a sexual cycle in the self-fertile heterokaryotic culture received from CBS815.71: the CBS815.71-sp3 strain with a mating-type allele orthologous to the *mat+* mating type of *Podospora anserina*, corresponding to the *MAT1-1* allele according to the official nomenclature (34), and the CBS815.71-sp6 strain with a mating-type allele orthologous to the *mat-* mating type of *P. anserina*, corresponding to the *MAT1-2* allele according to the official nomenclature (34). We assessed the frequency of second-division segregation (SDS) at the mating-type locus and, therefore, the frequency of heterokaryotic spore formation, by generating and isolating 190 four-spore asci from a cross between CBS815.71-sp3 and CBS815.71-sp6. We studied the size of the region in which recombination was fully abolished around the mating-type locus, by isolating 107 small ascospores from five-spore asci (monokaryotic) from a cross between CBS815.71-sp3 and CBS815.71-sp6. After isolation, the strains were maintained on M2 minimal medium to prevent contamination (71), and were incubated at 22°C for growth. The growth of CBS815.71-sp3 and CBS815.71-sp6 homokaryons and the CBS815.71 dikaryotic strain was assessed on OA, V8 and M2 media (71). The diameter of mycelial colonies was measured after 3 and 10 days of growth at 22°C with 12 h illumination, with a 10 cm ruler. Mean growth rate per day was calculated over the seven-day period from day 3 to day 10. Dry weights were measured after 10 days of growth in liquid M2 medium (50 mL); the mycelium was filtered and dried at 65°C overnight. Measurements were performed on five Petri dishes for each strain and each medium. ANOVA was performed with R software v 4.2.0.

### Genome sequencing and other genomic data

We generated long read-based genome assemblies by sequencing the genomes of the *S. tetrasporum* strains CBS815.71-sp3 and CBS815.71-sp6 with both Illumina technology on the HiSeq2500 platform (150 bp long paired-end reads) and Oxford Nanopore MinION technology with an R9 flow cell. For each DNA extraction, we prepared 100–200 mg aliquots of mycelium, which were stored in a 1.5 mL Eppendorf tube at −80°C for at least 24 hours before lyophilization. A sheet of cellophane was placed over the M2 medium before inoculation with spores, such that the mycelium grew on the cellophane for three to four days and could be scraped off easily. For Illumina sequencing, DNA was extracted with the commercial Nucleospin Soil kit from Macherey Nagel. Illumina library preparation and genome sequencing were outsourced to the sequencing core facility of Genoscope (last accessed May 15, 2021). For Nanopore sequencing, mycelium DNA was extracted with the NucleoBond High Molecular Weight DNA kit from Macherey-Nagel, with the mechanical disruption of about 30 mg of lyophilized mycelium with beads for 5 min at 30 Hz. The Nanopore library was prepared with the SQK-LSK109 ligation sequencing kit, and sequencing was performed in house. The quality of the raw reads was analyzed with fastQC 0.11.9 (72) for the Illumina raw reads and pycoQC 2.5.0.23 (73) for the Nanopore raw reads (Table S5).

In addition, the genome and transcriptome of *S. tetrasporum* strain CBS815.71-sp3 were sequenced with Illumina technology by the Joint Genome Institute (JGI). DNA was extracted following (74) from mycelium grown on cellophane-covered potato dextrose agar medium. Plate-based DNA library was prepared for Illumina sequencing on the PerkinElmer Sciclone NGS robotic liquid handling system with the Kapa Biosystems library preparation kit. Sample DNA (200 ng) was sheared into 600 bp fragments with a Covaris LE220 focused ultrasonicator. The sheared DNA fragments were size-selected by double-SPRI and the selected fragments were end-repaired, A-tailed, and ligated with Illumina-compatible sequencing adaptors from IDT containing a unique molecular index barcode. RNA was extracted from the same mycelial colonies as DNA using TRIzol (Invitrogen) following manufacturer’s instructions (https://assets.thermofisher.com/TFS-Assets/LSG/manuals/trizol_reagent.pdf). Plate-based RNA sample preparation was obtained with the PerkinElmer Sciclone NGS robotic liquid handling system and the Illumina TruSeq Stranded mRNA HT sample preparation kit, with the poly-A selection of mRNA according to the protocol in the Illumina user guide (https://support.illumina.com/sequencing/sequencing_kits/truseq-stranded-mrna.html), under the following conditions: 1 µg total RNA as the starting material, and eight PCR cycles for library amplification. The prepared libraries were then quantified with the KAPA Biosystem next-generation sequencing library qPCR kit (Roche) and a Roche LightCycler 480 real-time PCR instrument. The quantified libraries were then then multiplexed with other libraries, and the pool of libraries was prepared for sequencing on the Illumina NovaSeq 6000 sequencing platform with NovaSeq XP v1.5 reagent kits, an S4 flow cell, following a 2×150 indexed run method. All raw Illumina sequence data were filtered for artifacts/process contamination. DNA reads produced by the JGI were then assembled with SPAdes v3.15.2 (75). The obtained genome assembly is referred to as the Illumina read-based assembly of CBS815.71-sp3.

The genome of the *P. anserina* S *mat+* strain “Podan2” (32) was downloaded from the JGI MycoCosm website (https://mycocosm.jgi.doe.gov/mycocosm/home, last accessed November 15, 2018) and improved annotations by (76) were downloaded from the GitHub repository, https://github.com/johannessonlab/SpokBlockPaper (last accessed June 1, 2020). The genome of the OR74A *N. crassa mat A* strain (77) was downloaded from GenBank under accession number GCA_000182925.2 (last accessed May 10, 2022).

### Long read-based genome assemblies for CBS815.71 strains

*De novo* assemblies of the genomes of the CBS815.71-sp3 and CBS815.71-sp6 strains were constructed from both Illumina reads produced at Genoscope and Nanopore reads. For each genome, the raw Nanopore reads were trimmed and assembled with Canu v1.8 (78) with the option genomeSize=35m. The assembly obtained with Canu was polished twice with Nanopore trimmed reads using first Racon v1.4.3 (79) with the options m=8 x=6 g=8 w=500 and then with medaka (https://github.com/nanoporetech/medaka) with the default settings. The assembly obtained was polished five times with the trimmed Illumina reads with Pilon v1.24 (80). For each round of polishing, BWA v0.7.17 (81) was used to align the Nanopore or Illumina trimmed reads with the assembly for polishing. The Illumina reads were first trimmed with Trimmomatic v0.36 (82) with the options: ILLUMINACLIP:TruSeq3-PE.fa:2:30:10 LEADING:10 TRAILING:10 SLIDINGWINDOW:5:10

MINLEN:50. The quality of the final assemblies was assessed with Quast v5.1 (83). We checked for contamination, by performing a BLAST analysis of the assembly scaffolds against the online NCBI nucleotide collection database (https://blast.ncbi.nlm.nih.gov/, last accessed March 5, 2022) and looking at the coverage of Illumina and Nanopore reads on the final assemblies. We also evaluated the completeness of the final assemblies with BUSCO v5.2.2 (84,85) and the Sordariomyceta ortholog set (sordariomycetes_odb10, 2020-08-05, *n*= 3817 orthologous groups searched). We also ran BUSCO analysis on the Podan2 *P. anserina* assembly for comparison.

### Genome annotation

Before gene annotation, the final assemblies were formatted (contigs were renamed and ordered in descending order of size) and masked for repeats with RepeatMasker (86) and the funnannotate v1.8.9 pipeline (87). We performed gene annotation on the masked assemblies with BRAKER v2.1.6, which uses a combination of the *ab initio* gene predictors Augustus and GeneMark-ET (88). We ran BRAKER twice, first using the BUSCO dataset of proteins sordariomycetes_odb10 (84) as evidence, and then using the RNAseq dataset as evidence. We optimized the training steps, by concatenating the two homokaryotic genomes derived from the same heterokaryotic strain, CBS815.71, into a single fasta file and running BRAKER on the combined fasta file. For RNAseq mapping, we used STAR v2.7.10a (89) with the default settings. We combined the results of the two BRAKER runs with TSEBRA (the Transcript Selector for BRAKER; https://github.com/Gaius-Augustus/TSEBRA), with the option ‘intron_support’ set to 0.2. We ran InterProScan5 v5.54-87.0 (90), TMHMM v2.0c (91), SignalP v4.1 (92) and Phobius (93) and obtained predicted protein annotations from PFAM, InterPro, EggNog, UniProtKB, MEROPS, CAZyme, and GO ontology. For repeat annotation, we performed a *de novo* identification of transposable elements in the final assemblies with RepeatModeler v1.0.11 (94). We merged the consensus sequences obtained wiereth the RepBase library available in RepeatMasker v4.0.9 (86) and the custom library “PodoTE-1.00” built for the *Podospora anserina* species complex by (95) and the *Neurospora* library of (96) “Gioti_neurospora_repeats.renamed” (both available from the GitHub repository https://github.com/johannessonlab/SpokBlockPaper, last accessed June 1, 2020). We used RepeatMasker v4.0.9 (86) to annotate repeats. We parsed the RepeatMasker outputs and removed low-complexity and simple repeats with the parseRM_merge_interrupted.pl script from https://github.com/4ureliek/Parsing-RepeatMasker-Outputs (last accessed February 25, 2021). We retained only TEs longer than 150 bp, and overlapping TEs annotated as belonging to the same family were merged into single elements for the final annotation. We used the TE annotation suggested by (97) to sort TEs by family. We looked for signatures of RIP with the RIPper platform http://theripper.hawk.rocks (default settings; (98)). The mating-type locus was identified by tBLASTn, with the *mat+* and *mat-* idiomorphs of *P. anserina* as queries. The *SMR1, SMR2, FMR1, FPR1* gene models were manually curated as previously described (99), in Artemis v18.0.2 (100). The sequences of proteins containing a MATα_HMG domain were downloaded from the European Nucleotide Archive (ENA) database with accession numbers CDN29981 and ESA43845 for *P. anserina* and *N. crassa*, respectively. Proteins were aligned with Clustal Omega (101) and the alignments were color-coded with JALVIEW v2 (102).

### Comparative genomics and statistical analyses

For all comparative genomics analyses, we identified orthologous groups with OrthoFinder v2.3.7 (103). We calculated dS values by studying one-to-one orthologs between the CBS815.71-sp3 and CBS815.71-sp6 genome assemblies and retaining only those orthologs with the same number of exons and the same transcript length as predicted. We aligned orthologous gene coding sequences, using the codon-based approach implemented in translatorX v1.1 with default parameters (104). We used the nucleotide alignment as input for the yn00 program in the PAML v.4.9 package (105,106). Pairwise Wilcoxon tests were performed in R with false discovery rate correction. We excluded the four genes located at the edges of strata to assess the differences in dS between strata. We studied synteny between assemblies, by investigating either collinearity between one-to-one orthologs or running the genome sequence aligner nucmer from MUMmer v4.0.0rc1 (107) with default options. For the identification of small indels between the long-read assemblies, we first aligned the CBS815.71-sp6 genome assembly with the CBS815.71-sp3 genome assembly using Minimap2 (108) with the -cx asm10 -t8 options. We then used samtools mpileup (109) with the following options: -d 250 -m 1 -E -- BCF --output-tags DP,DV,DP4,SP.

### Genotyping of the mating-type locus alleles and surrounding regions

We devised a new method of DNA extraction for genotyping based on a published protocol (110). Briefly, 1 mm^3^ plugs were taken directly from cultures growing on agar plates and ground with 200 µL extraction buffer, heated at 95°C for 15 minutes, placed on ice for one minute, briefly vortexed and centrifuged at 15,900 x g for 30 seconds. We determined the mating-type allele of the offspring from a cross between CBS815.71-sp3 and CBS815.71-sp6 by PCR with the primers and conditions described in Table S6 and Table S7. Primers were designed with Primer3 software (111,112), based on the mating-type locus region annotated in the two genome assemblies obtained for *S. tetrasporum*. We visualized amplicons by agarose-gel electrophoresis.

## Data availability

All genomic data produced by the JGI are available from the JGI MycoCosm website (https://mycocosm.jgi.doe.gov/mycocosm/home, last accessed February 5, 2022). The raw Illumina and Nanopore reads and the long read-based assemblies for the CBS815.71-sp3 and CBS815.71-sp6 genomes were deposited in GenBank under the BioProject accession number PRJNAXXX (BioSample ID: xxx, assembly accession number: xxx; available upon publication).

## Acknowledgements

This work was supported by the Louis D. Foundation award and EvolSexChrom ERC advanced grant #832352 to T.G., DOE-JGI CSP grant #504394 to P.G. and P.S, the work (proposal 10.46936/10.25585/60001199) conducted by the US Department of Energy Joint Genome Institute (JGI : https://ror.org/04xm1d337), a DOE Office of Science User Facility, and supported by the Office of Science of the US Department of Energy under contract no. DE-AC02-05CH11231. We are grateful to J.P. Vernadet for sharing the conda environment to run the BRAKER pipeline.

## Author contributions

T.G., F.E.H. and P.S. conceptualized the study, acquired funding and supervised the study. N.V., A.S., V.G., E.D.F., E.L., F.E.H., and P.S. contributed to experiments. N.V., F.E.H., R.D., C.L., P.S. and R.R.d.l.V. analyzed the genomes. N.V., V.G., E.D.F., E.L. and P.S. analyzed the progenies. P.G., S.G., Y. Z., S. T. and I.G. provided RNAseq data and the Illumina based-genome assembly of the CBS815.71-sp3 strains. N.V., F.E.H., P.S., and T.G. wrote the manuscript draft. All authors edited the manuscript.

## Supporting information captions

**Supplementary note S1:** Frequency of second-division segregation of the mating-type locus under random segregation of the mating-type locus.

**Supplementary Table S1:** Tests of differences in growth speed and dry weight between *Schizothecium tetrasporum MAT1-1 and MAT1-2* homokaryons and heterokaryons on different media.

**Supplementary Table S2:** Genome assembly of the *Schizothecium tetrasporum* monokaryotic strains CBS815.71-sp3 and CBS815.71-sp6.

**Supplementary Table S3:** List of 15 genes in the recombination suppression region around the mating-type locus in *Schizothecium tetraporum* predicted to encode proteins that are putative transcription factors or involved in signalling pathways.

**Supplementary Table S4:** List of genes in the recombination suppression region around the mating-type locus in *Schizothecium tetraporum* having orthologues also located in the recombination suppression region around the mating-type locus in *Podospora anserina* and *Neurospora tetrasperma*.

**Supplementary Table S5:** Quality of the raw reads produced and used in this study.

**Supplementary Table S6:** PCR primers used in this study.

**Supplementary Table S7:** PCR conditions used in this study.

**Supplementary Figure S1:** Genome collinearity between the long read-based assembly and the Illumina-based assembly of *Schizothecium tetrasporum* CBS815.71-sp3 strain.

**Supplementary Figure S2:** Transposable element annotation in different genomic regions of the *Schizothecium tetrasporum* CBS815.71 strain.

**Supplementary Figure S3:** Amino-acid alignment of proteins containing a MATα_HMG domain.

**Supplementary Figure S4:** Genome-wide distribution of synonymous divergence (dS) in the *Schizothecium tetrasporum* CBS815.71 strain. dS was computed per-gene between the CBS815.71-sp3 and CBS815.71-sp6 assemblies and plotted according to the CBS815.71-sp3 assembly gene position.

**Supplementary Figure S5:** GC content variation and determination of centromeres the *Schizothecium tetrasporum* CBS815.71-sp3 assembly.

**Supplementary Figure S6:** Delimitation of three evolutionary strata in the S*chizothecium tetrasporum* CBS815.71 strain.

**Supplementary Figure S7:** Probability of recombination in centimorgans (cM) between markers and the mating-type locus in the heterokaryotic strain CBS815.71 after one round of experimental selfing measured in 107 of the offspring.

**Supplementary figure S8:** Genome-wide collinearity based on single orthologous genes between *Neurospora crassa, Podospora anserina* and *Schizothecium tetrasporum* genomes.

## Notes

### Competing Interest Statement

The authors have declared no competing interest.

### Summary of Updates

Author affiliation information

